# Caterpillar counter-adaptations to hooked trichomes: does the coevolutionary arms race continue?

**DOI:** 10.1101/2025.03.22.644295

**Authors:** Ritabrata Chowdhury, Erika C.P. de Castro, Glennis Julian, Simon Chen, Chris D. Jiggins, Walter Federle

## Abstract

Plants protect themselves against herbivory with diverse chemical and physical defences, and many herbivores have evolved counter-adaptations against these defences. So far, little is known about herbivore counter-adaptations to physical plant defences and their underlying mechanisms. Here, we investigated how specialised Heliconiini caterpillars are able to cope with the hooked trichomes of *Passiflora adenopoda* vines. *P. adenopoda* trichomes have sharp tips reinforced with silica, in contrast to simpler trichomes of other *Passiflora* that do not deter Heliconiini larvae. While *Heliconius melpomene* and *H. erato* caterpillars placed on *P. adenopoda* leaves were arrested and their soft cuticle was pierced by the sharp hooked trichomes, *H. charithonia* and *Dryas iulia* caterpillars of all sizes could move easily over the leaves without any injury. The superior performance of these two species on trichome-bearing leaves can be explained by their thicker and more puncture-resistant cuticle. Morphological and penetrometry measurements showed that the relevant cuticle regions in *H. charithonia* and *D. iulia* caterpillars are significantly thicker and more puncture-resistant than in *H. melpomene* and *H. erato*. When we tested the ability of the caterpillars to feed on *P. adenopoda* leaves with hooked trichomes, only *H. charithonia* survived, indicating the existence of further, previously unknown post-ingestive adaptations. Understanding insect counter-adaptations to physical plant defences is essential for studying the evolutionary arms-race between plants and insect herbivores.

## Introduction

Plants and insects have lived together for more than 400 million years and have evolved complex interactions and adaptations to one another (1–3). While many relationships are mutually beneficial, such as pollination or seed dispersal, most interactions involve insects feeding on plants, and plants defending themselves (3). It has been proposed that insect-plant interactions and the adaptations arising from them are a key factor in promoting the diversity of both herbivores and host plants (1, 4). Insect-plant interactions are mediated by chemical, physical, and/or behavioural traits. While the chemical ecology of these interactions has been studied in some detail (5–9), the role of physical factors is less well explored (10, 11). In particular, little is known about how herbivores have adapted to physical plant defences. The co-evolution of insects and chemical plant defences has been described as an arms-race (1, 4, 12), but it is unclear if physical plant defences can have a similar outcome.

We investigate here the role of hooked trichomes in *Passiflora* vines as a defence against Heliconiini (Nymphalidae: Heliconiinae) caterpillars.The Heliconiini tribe is a diverse group of butterflies found mainly in South and Central America that feed on *Passiflora* and closely related genera during their larval stages (13). *Passiflora* and their Heliconiini herbivores show mutual adaptations providing a well-studied example of co-evolution and adaptive radiation (14–16). *Passiflora* have evolved multiple chemical and biotic anti-herbivory defences including toxic compounds such as cyanogenic glycosides, alkaloids, tannins, and phenolics (17), diverse leaf shapes and variegation (to disguise their identity) (16, 18), leaf structures mimicking butterfly eggs to deter oviposition (16, 19), as well as extrafloral nectaries to attract ants (20). On the other hand, Heliconiini have adapted by developing highly accurate visual and chemosensory systems (21, 22), behavioural strategies to overcome *Passiflora* defences (14, 23), and the ability to sequester the cyanogenic glycosides of their *Passiflora* hosts for their own defence (17, 24, 25). In addition to chemical and biotic defences, several *Passiflora* have evolved trichomes as a physical defence against herbivores (26–29).

Trichomes are hair-like structures of epidermal origin that can play a role in herbivore defence (30) (31, 32). While some glandular trichomes release toxic or sticky substances (33, 34), non-glandular trichomes can enhance plant defence by physically inhibiting herbivore movement, limiting herbivore access to leaf or stem tissues (35–37), or reducing insect oviposition (38, 39). In response, insect herbivores have adapted to handle plant trichomes. Some of the few known adaptations are morphological, such as elongated legs to avoid contact with the sticky heads of glandular trichomes or tarsal hooks to anchor onto trichomes (40). Others involve behaviours such as bugs walking "on tiptoe" between trichomes (41), caterpillars removing trichomes before feeding on the underlying tissues (42, 43) or caterpillars spinning silk mats to move on trichome-bearing surfaces without being punctured (44).

Some *Passiflora* species (*P. lobata, P. adenopoda*) possess stiff, hook-shaped trichomes with a sharp tip on their leaves. These trichomes can injure caterpillars by piercing their cuticle, and kill them due to a combination of starvation and blood loss (45, 46). Similar dramatic effects have been observed in *Phaseolus* beans, where hooked trichomes trap aphids and bed bugs by piercing the cuticle in different parts of the legs (47–51) and in an ant-plant, *Macaranga trachyphylla*, where hooked trichomes pierce and kill lycaenid *Arhopala* caterpillars (52). It has been suggested that by evolving hooked trichomes, *P. adenopoda* has won the co-evolutionary arms race with its specialised Heliconiini herbivores (45). However, some Heliconiini are not deterred even by these defences. Among the ca 75 known species of Heliconiini, four have been reported to be able to walk on *Passiflora* with hooked trichomes without being injured: *Dione moneta* (14) *Heliconius charithonia*, *Agraulis vanillae* and *Dryas iulia* (46). However, the exact mechanism by which the caterpillars accomplish this feat is still unclear, as are the adaptations involved. Cardoso (46) suggested that the caterpillars can cope with hooked trichomes by laying silk mats over them, biting off trichome tips, or forcefully pulling out their prolegs to avoid entrapment; he also speculated that adaptations of the cuticle may be involved. However, these hypothetical mechanisms have not been studied in detail and it is not known whether any morphological adaptations of the cuticle and prolegs are involved. Surprisingly, only *H. charithonia* is known to feed on *P. adenopoda* under natural and insectary conditions (53, 54). It is unclear whether any other species can do this and whether the hooked trichomes have any post-ingestive effect.

Here, we investigate the mechanisms that enable specialised Heliconiini caterpillars to cope with the hooked trichomes of *P. adenopoda*, and whether the hooked trichomes have a post-ingestive effect on the caterpillars.

## Results

### Morphology of hooked trichomes in *P. adenopoda*

The leaf and stem surfaces of *P. adenopoda* are densely covered by non-glandular, unbranched, hook-shaped trichomes (Figs. 1.A & 1.B). Their density was higher on the underside (abaxial side) than on the upper (adaxial) side of the leaf (3.5+/-0.67 vs. 1.91+/-0.20 trichomes/mm²; mean +/- SD, n=5 leaves). The hooked trichomes of *P. adenopoda* occur in different sizes on both sides of the leaves and ranged in length from 50 to 430 µm (Fig.S1). The trichomes are very sharp (tip radius 0.51+/-0.19 µm; mean +/- SD, n=5 trichomes from 3 different leaves). EDX elemental mapping images from *P. adenopoda* showed a high concentration of silicon (Si) (colour-coded green, Figs 1.C) at the base and tip of hooked trichomes, whereas in between, there is a strong deposition of calcium (Ca) (colour-coded blue, Fig 1.C).

**Fig. 1.**
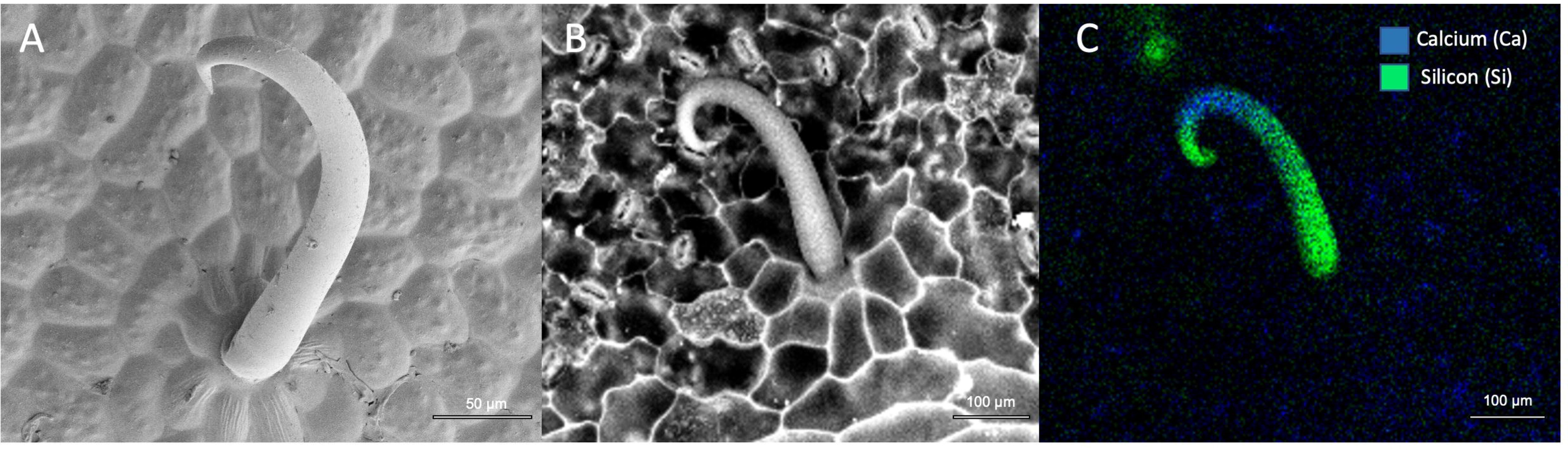
(*A*) Electron micrograph of a *P. adenopoda* trichome from the upper side of a leaf; (*B*) and (*C*) Backscattered-electron image (detecting differences in atomic number) and corresponding EDX image of a *P. adenopoda* trichome from the underside of a leaf, colour-coded for the presence of silicon (green) and calcium (Ca).

### Hooked trichomes arrest the movements of *H. melpomene* and *H. erato* caterpillars by piercing their cuticle

When *H. melpomene* and *H. erato* caterpillars were placed on *P. adenopoda* leaves, they were unable to move more than a few mm, because they were arrested by one or several hooked trichomes. This had a clear impact on walking speeds: *D. iulia* was the fastest, followed by *H. charithonia,* and whereas *H. erato* and *H. melpomene* were effectively unable to move (Fig. 2.A) (one-way ANOVA; F_3,32_=21.63, p <0.001). However, when caterpillars were tested on "shaved" *P. adenopoda* leaves, walking velocities significantly increased for *H. erato* and *H. melpomene,* but not for *H. charithonia* and *D. iulia* (Fig 2A, Table S1). Indeed, *H. erato* and *H. melpomene* were as fast as *D. iulia* on “shaved” leaves of *P. adenopoda* (Fig.2A).

**Fig. 2.**
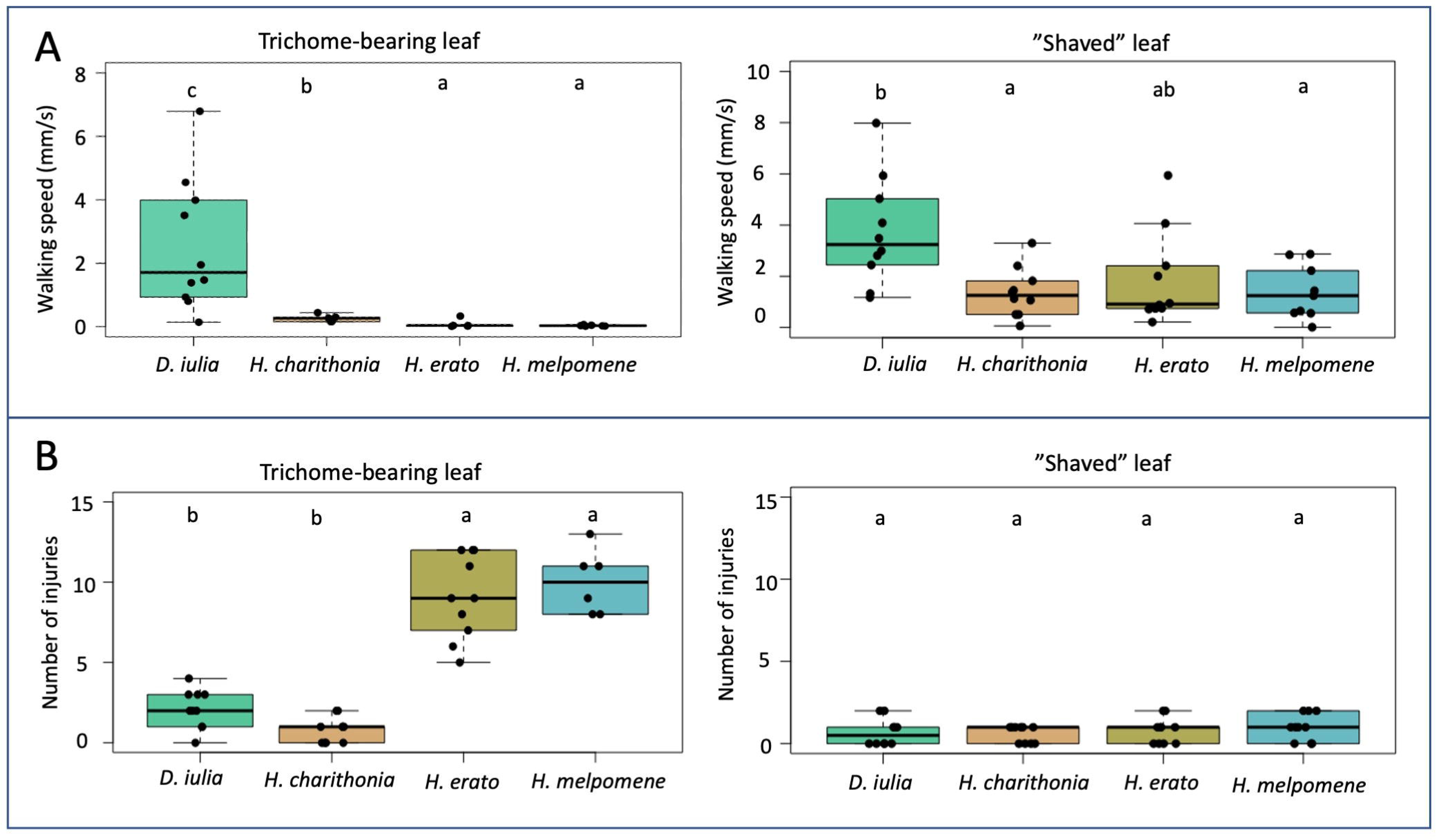
(*A*) Walking velocity of different caterpillars on *P. adenopoda* leaves with intact trichomes vs. "shaved" leaves; (*B*) Number of injuries sustained by different caterpillars on *P. adenopoda* leaves with intact trichomes vs. "shaved" leaves; in the box plots, different letters indicate statistically significant differences between groups according to Tukey’s HSD or Dunn’s test (p < 0.05), while groups sharing the same letter are not significantly different.

Video recordings of small 1^st^-2^nd^ instar caterpillars on *P. adenopoda* leaves showed that the tips of the hooked trichomes touched or pierced their body wall at about half the caterpillar’s body height. Piercing of the cuticle of small caterpillars was observed for *H. erato* and *H. melpomene* (see Movie S1), but not for *H. charithonia* and *D. iulia*. In larger 3^rd^-4^th^ instar caterpillars, it was also found that only *H. erato* and *H. melpomene* were pierced but not *H. charithonia* and *D. iulia*. However, the main area where the trichomes pierced the cuticle was the soft distal planta of the prolegs (including the corona, subcorona and smooth pad) (55, 56), sometimes leading to bleeding (Figs. 3A, B, C; Movie S2). The trichomes caused injuries at multiple locations on the planta (evident from methylene blue staining or directly visible in SEM; Figs. 3D, E, F). However, in the case of *H. charithonia* and *D. iulia*, the trichomes did not penetrate the cuticle. If the trichomes briefly interlocked, the caterpillars were able to escape the entrapment by pulling away their prolegs. Virtually no injuries were observed in *D. iulia* and *H. charithonia* caterpillars (see Movies S3 & S4).

**Fig. 3.**
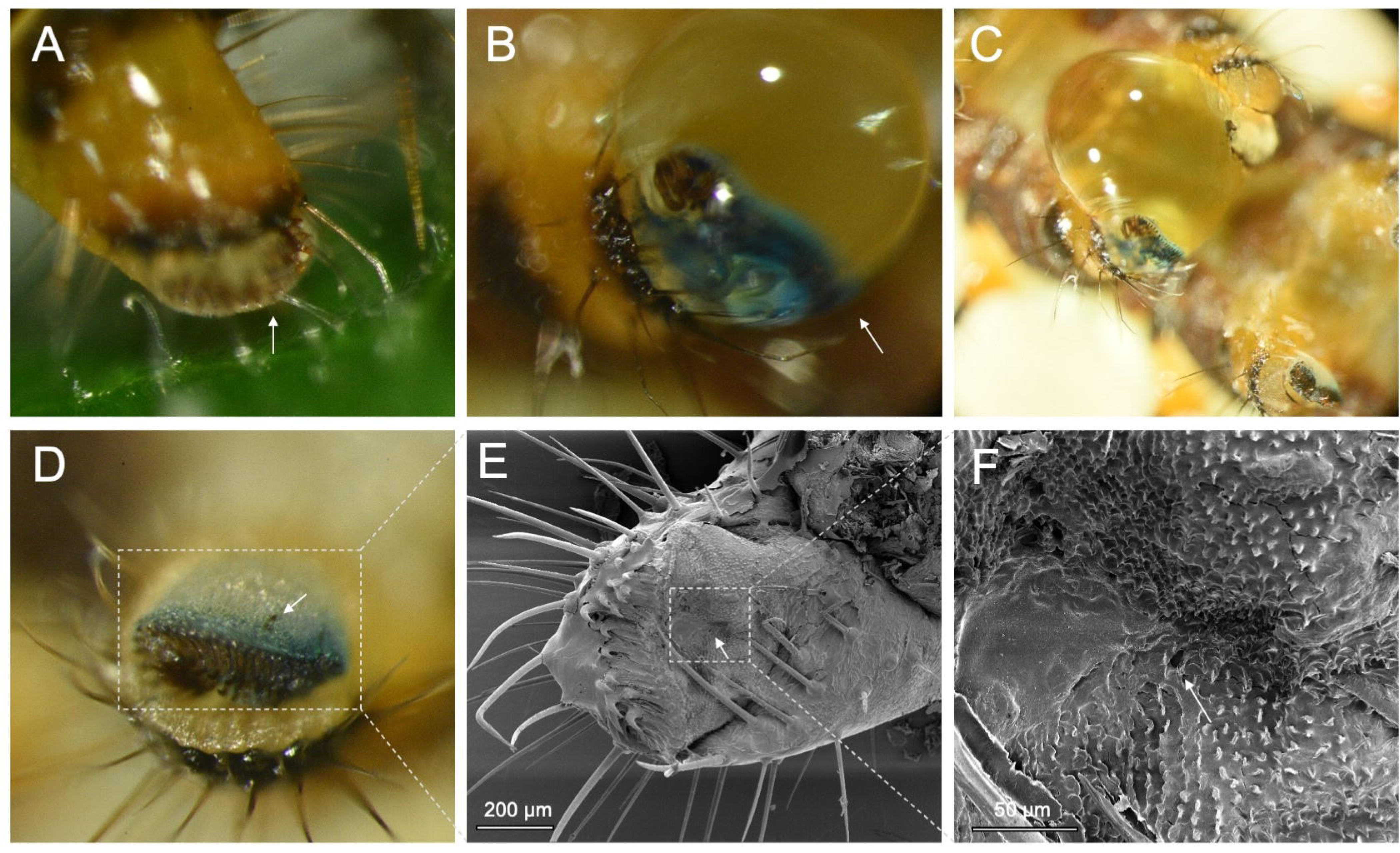
(*A*) Hooked trichome pierced into the soft planta of the proleg of a later instar *H. erato* caterpillar (arrow); (B) and (C) Haemolymph (arrow) oozing from the wounds on the footpad of a later instar *H. erato* caterpillar after interacting with the hooked trichomes; (*D*) Footpad injury of a later instar *H. erato* caterpillar as observed after methylene blue staining; one injury spot shown by arrow; (*E*) and (*F*) SEM images of the same trichome-inflicted injury.

The number of injuries on the prolegs after walking on *P. adenopoda* leaves with trichomes differed significantly between caterpillar species (Kruskal-Wallis test; c² = 28.1, df= 3, p<0.001) (Fig. 2.B). *H. melpomene* and *H. erato* had significantly more injuries than *H. charithonia* and *D. iulia* (Table S2, Fig. 2.B). SEM images confirmed that the methylene blue staining marks corresponded to injuries inflicted by the sharp, hooked trichomes (Figs 3.D, E & F). When placed on "shaved" *P. adenopoda* leaves, the number of injuries was drastically decreased in *H. erato* (Wilcoxon signed rank test; n=10, W= 100, p<0.01) and *H. melpomene* (Wilcoxon signed rank test; n=10, W= 60, p<0.01) whereas the numbers remained low for *H. charithonia* (Wilcoxon signed rank test; n=10, W=56, p>0.05) and *D. iulia* (Wilcoxon signed rank test; n=10, W=73, p>0.05).

### Caterpillar species differ in puncture resistance and cuticle thickness

The puncture resistance of the body cuticle and the prolegs varied significantly between the caterpillar species and across different sizes/instars (Fig 4A, B & C). In the smallest (1^st^ and 2^nd^ instar) larvae, both *H. charithonia* and *D. iulia* caterpillars showed a higher puncture resistance of the body cuticle than *H. erato* and *H. melpomene* (Fig. 4A, one-way ANOVA, F_3,32_=8.67, p<0.001, Table S3). Similarly, when puncture resistance was measured in the prolegs of 3^rd^-4^th^ instar caterpillars, *D. iulia* and *H. charithonia* had significantly higher values than *H. erato* and *H. melpomene* (Fig. 4.B, Table S4). A different result was found for the body cuticle of large (3^rd^-4^th^ instar) caterpillars (one-way ANOVA, F_3,36_=9.47, p<0.001), where *D. iulia* was more puncture-resistant than *H. erato*, *H. melpomene* and *H. charithonia* (all Tukey HSD p<0.01) (Fig. 4.C, Table S5).

**Fig. 4.**
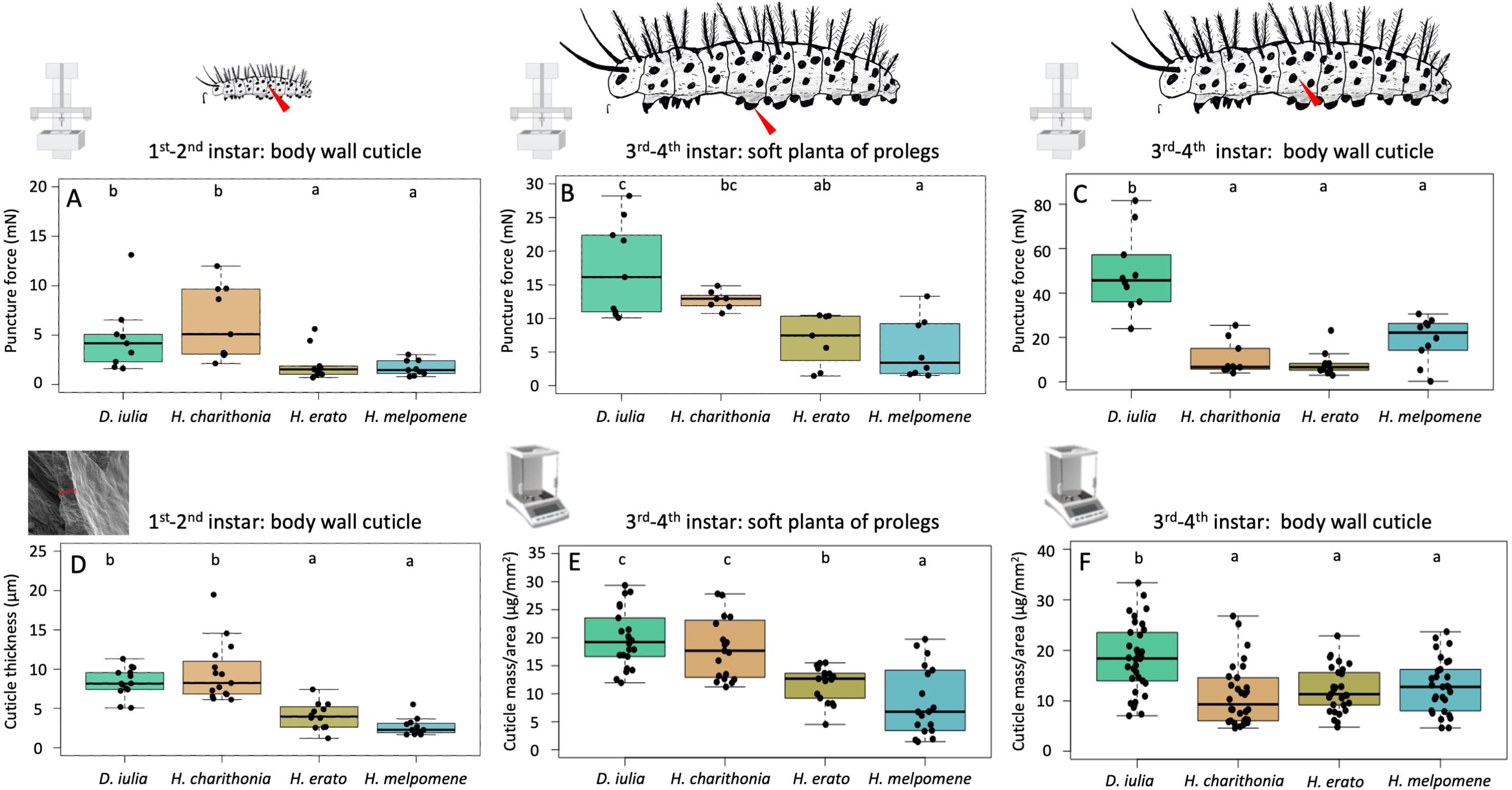
Puncture resistance, measured as the force required to puncture different cuticle regions of caterpillars of four different Heliconiini species. (*A*) Body wall cuticle of small (1^st^-2^nd^ instar) caterpillars; (*B*) Soft planta of the prolegs in large (3^rd^-4^th^ instar) caterpillars; (*C*) Lateral abdominal cuticle of large (3^rd^-4^th^ instar) caterpillars; (*D*) Cuticle thickness of body wall in small (1^st^-2^nd^ instar) caterpillars, and; (*E*) and (*F*) Area-specific mass of the soft planta of prolegs in large (3^rd^-4^th^ instar) caterpillars, and of the lateral abdominal cuticle of large (3^rd^-4^th^ instar) caterpillars, respectively; in the box plots, different letters indicate statistically significant differences between groups according to Tukey’s HSD test (p < 0.05), while groups sharing the same letter are not significantly different

The differences in puncture resistance between caterpillar species are consistent with measurements of cuticle thickness. The thickness and area-specific dry mass of the lateral body wall and proleg cuticle closely followed the observed species differences in puncture resistance, for both small and large caterpillars (Figs 4.D, E & F, Tables S6,S7,S8). As the measurement of area-specific dry mass for the smallest caterpillars had a relative error that was too large, we measured cuticle thickness from freeze-fractured cuticle samples. The data confirmed that the body wall cuticle of 1^st^-2^nd^ instar caterpillars, and the soft planta of the prolegs of 3^rd^-4^th^ instar caterpillars were significantly thicker in *D. iulia* and *H. charithonia* than in *H. melpomene* and *H. erato* (Fig 4.D,E, Table S6, S8), whereas for the body wall cuticles of 3^rd^-4^th^ caterpillars, *D. iulia* had significantly higher values than the other three species (Fig 4.F, Table S7).

### Only *H. charithonia* caterpillars can successfully develop on *P. adenopoda*

We investigated the ability of the four Heliconiini species (*H. charithonia, H. melpomene, H. erato* and *D. iulia*) to feed and develop on *P. adenopoda* (with intact trichomes) and trichome-free *P. biflora*. We found that only *H. charithonia* caterpillars could survive and successfully pupate on *P. adenopoda* shoots, whereas *H. erato*, *H. melpomene*, and *D. iulia* all died at an early stage (Table 1). Early-instar *H. erato* and *H. melpomene* fed on the leaves with hooked trichomes, but they were often impaled by trichomes before they could start feeding, and none of the larvae survived past the first or early second instar. Early-instar *D. iulia* caterpillars also fed on *P. adenopoda* and died quickly; only four larvae survived up to the early third instar. Dead larvae of *H. melpomene, H. erato* and *D. iulia* were either found near the leaf areas where they had been eating, suggesting that they died while eating, or in a dried-up state on the stem or leaves. In contrast, all four species successfully reached pupation when raised on *P. biflora* shoots (Table 1) and interestingly, survival of *H. charithonia* larvae was significantly better on *P. biflora* than on *P. adenopoda* (Fisher’s Exact test, p<0.001).

**Table 1.**
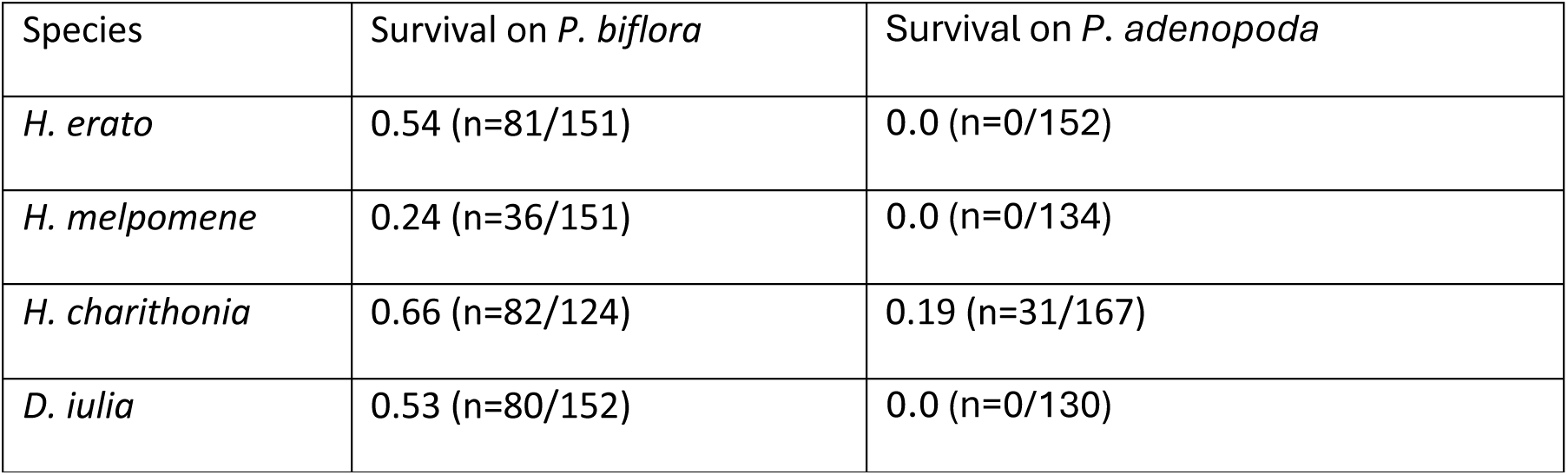
Larval survival (number of larvae that reached pupation per number of eggs from for each plant species) on *P. adenopoda* vs *P. biflora*.

## Discussion

Plant physical defences are wide-spread and form the first line of defence that insect herbivores have to overcome (3, 6). So far, little is known about the effects of plant physical defences on insect herbivores, and only few studies mention insect counter-adaptations against them (40–44). Our study shows that the hooked trichomes of *P. adenopoda* have two fundamentally different lethal effects on herbivorous caterpillars, one external when they walk over the leaves and the other internal when they eat the leaves. We also discovered that the ability of some specialized heliconiine caterpillars to walk on *Passiflora* species with hooked trichomes is associated with their thicker and more puncture-resistant cuticle.

### External effects of hooked trichomes on Heliconiini caterpillars

*Heliconius melpomene* and *H. erato* caterpillars placed on leaves of *P. adenopoda* were impaled by hooked trichomes and became immobilised, whereas *Dryas iulia* and *H. charithonia* caterpillars were able to walk easily across the leaves. As a result of multiple injuries caused by the trichomes, most *H. melpomene* and *H. erato* caterpillars lost haemolymph and all caterpillars died by blood loss, starvation and/or desiccation, whereas *D. iulia* and *H. charithonia* caterpillars remained effectively unharmed. The dramatic effects were clearly caused by the hooked trichomes of *P. adenopoda*, because all caterpillars were able to walk easily on the leaves and suffered no injuries when the trichomes were experimentally removed.

Our observations confirm earlier observations on large caterpillars (45, 46), but we found that small caterpillars are damaged in a different way. While in large caterpillars the trichome tips pierced the soft distal part of the abdominal prolegs (45), in smaller caterpillars (1^st^-2^nd^ instar) they caused injuries more dorsally on the lateral body wall, as the trichomes are larger relative to body size. The observed variation in trichome height within the same leaves may enhance the effectiveness of the physical defence, as the trichomes can reach a larger area of the herbivore’s body. The effect of trichomes on the youngest herbivores is particularly important. Whether an egg laid on a host plant can develop successfully into a pupa depends to a large extent on whether the newly hatched caterpillars can move around on the plant and start feeding (57). It is therefore significant that *D. iulia* and *H. charithonia* caterpillars showed a better ability to walk over hooked trichomes both as small and large caterpillars.

The hook-shape of the trichomes of *P. adenopoda* makes it difficult for caterpillars to pull out their body or prolegs once caught by a trichome; this would be much easier if the trichomes were straight (45, 47, 51). A further functionally important aspect of the morphology of *P. adenopoda* trichomes is their biomineralization. The high concentrations of Si at the tip and base of the trichome, and of Ca in the middle, likely increase the trichome’s stiffness and the hardness of its sharp tip (49, 58). This biomineralization is probably important to stabilize the trichomes during their interaction with herbivores and to prevent them from breaking off.

### What mechanisms allow *D. iulia* and *H. charithonia* caterpillars of different size to walk on leaves with hooked trichomes?

Cardoso (46) suggested that the caterpillars protect themselves by biting off trichome tips and laying silk mats. The caterpillars indeed showed both of these behaviours, but it is unlikely that they play an important role for the immunity of *D. iulia* and *H. charithonia* caterpillars to hooked trichomes. Firstly, we did not observe *D. iulia* and *H. charithonia* caterpillars biting off the tips of trichomes, and both species often walked over leaf sections with completely intact trichomes without being affected by them. The trichome tips observed by Cardoso (46) in the frass of *H. charithonia* and interpreted as evidence of biting could also represent partially digested, whole trichomes, as these have a higher silica content at the tip. Secondly, while we observed all four caterpillar species to lay silk when placed on *P. adenopoda* leaves, these thin silk threads did not provide any protection against the hooked trichomes. Most caterpillars lay silk threads on leaf surfaces to improve their attachment (59, 60). *D. iulia* and *H. charithonia* had no difficulty walking on *P. adenopoda* leaves even immediately after being placed on them, when the leaves were still devoid of silk threads.

Instead, the ability of *D. iulia* and *H. charithonia* to cope with hooked trichomes can be explained by their thicker and more puncture-resistant cuticle. When caterpillars of both species walked over leaves with trichomes, they frequently touched the hooked trichomes. However, the trichome tips did not penetrate their cuticle and they could therefore easily free themselves again. We found that in the prolegs of 3^rd^-4^th^ instar *D. iulia* and *H. charithonia*, the soft planta of the prolegs was not only significantly thicker but also more puncture-resistant than in *H. melpomene* and *H. erato*. We found the same effect for the body wall cuticle of 1^st^-2^nd^ instar caterpillars; again the cuticle of *D. iulia* and *H. charithonia* was both thicker and more puncture-resistant. In both cases, the more puncture-resistant cuticle can explain how *D. iulia* and *H. charithonia* caterpillars are insensitive to the hooked trichomes.

Interestingly, we did not find the same effect for the body wall of larger caterpillars: here, the cuticle of *H. charithonia* was as thin and susceptible to puncturing as that of *H. melpomene* and *H. erato* (Figs. 4.C, F). However, as the body wall of larger caterpillars does not come into contact with the hooked trichomes, this does not play a role for *H. charithonia*. The local strengthening of specific cuticle regions in *H. charithonia* suggests that it has evolved as an adaptation to hooked trichomes, in contrast to *D. iulia*, where the caterpillars have a generally more robust body and thicker cuticle, as well as a wider host plant range (14).

A high cuticle puncture resistance is not only important for insects resisting sharp plant trichomes as we found in this study, but also for insects that defend themselves against predators and intraspecific opponents. Puncture resistance in such systems was also found to increase with cuticle thickness, but it is likely that material properties and sclerotization also play a role (61–63).

### Evidence for internal, post-ingestive effects of hooked trichomes on Heliconiini caterpillars

Previous studies on hooked trichomes have considered almost exclusively their external effects on insect herbivores (31, 45, 46, 50, 51, 64, 65). However, it is clear that to use trichome-bearing plants as hosts, herbivorous insects must not only be able to walk over their surfaces, but also feed on their leaves, and the sharp trichomes move through their digestive system. One possible consequence of the biomineralization of hooked trichomes is that their sharp tips pass undigested through the gut of insect herbivores (66, 67), making it more likely that they pierce the gut wall. We discovered that of the four species studied, the caterpillars of *H. charithonia* were the only ones able to feed and survive on *P. adenopoda* with hooked trichomes. *D. iulia* caterpillars could walk rapidly across *P. adenopoda* leaves, but were unable to feed and develop on *P. adenopoda*; their larvae attempted to feed on the leaves, but died quickly and only survived up to the third instar. This shows that *H. charithonia* caterpillars have both adaptations of their cuticle to externally resist the hooked trichomes and internal adaptations to cope with the trichomes in their digestive system. Post-ingestive effects of trichomes are probably widespread in larval and adult herbivores (68, 69), but so far the underlying mechanisms have not been studied in detail, and nothing is yet known about counter-adaptations of insects.

## Conclusion

While physical defences are widespread in plants (6, 45, 49, 70), our study is one of the first to characterise insect counter-adaptations. We show that *H. charithonia* and *D. iulia* caterpillars have a more puncture-resistant and robust cuticle, allowing them to resist the sharp, hooked trichomes of *P. adenopoda*, which can be fatal to caterpillars of other species. Only *H. charithonia* caterpillars were able to feed and survive on *P. adenopoda*, demonstrating that both external reinforcement of the cuticle and internal adaptations are responsible for its ability to handle the hooked trichomes. External and internal counter-adaptations of herbivores to physical plant defences such as the ones found in this study may be widespread but remain poorly documented.

## Materials and Methods

### Rearing of *Passiflora* plants and Heliconiini butterflies

*Passiflora adenopoda* (subgenus *Decaloba*, Section *Pseudodysosmia*) were grown from seeds (Plant World Ltd., Devon, UK, and Rareplantseed Ltd., München, Germany) in plastic pots (7×7×7cm) and plants were kept at 24-28 °C, 60-80% humidity and 12h day/night in the University of Cambridge greenhouses at Madingley. *P. biflora* Lam. (subgenus *Decaloba*) (subgenus *Passiflora*) plants were from *Passiflora* stocks originally obtained from Panama.

Four Heliconiini species were used in this study: *Heliconius charithonia*, *Heliconius erato*, *Heliconius melpomene*, and *Dryas iulia*. *H. erato* were from collected from the wild in Gamboa, Panama, in 2015, and subsequently raised and maintained as stocks in Finland and in Cambridge since 2017. *H. melpomene* stocks were from Stratford Butterfly Farm Ltd (Stratford-upon-Avon, UK). Populations of *H. charithonia* and *D. iulia* were established from pupae (∼40 per species) ordered from The Entomologist Ltd (East Sussex, UK).

Eggs were collected weekly from breeding cages and raised in larval cages in a climate-controlled room (24-28 °C, 80% RH and 12h day/night). Larval cages were supplied *ad libitum* with fresh *Passiflora* cuttings and checked every other day for pupae.

*H. charithonia* eggs were collected and larvae were raised on either *P. biflora* or *P. adenopoda* plants/cuttings. Pupae were hung in cages (40 x 40 x 60 cm) from a microfiber cloth wrapped around a bamboo stick. Pupal cages were kept in a climate-controlled room at 24-28 °C, 80% RH and 12h day/night. Pupal cages were checked every other day and recently emerged adult butterflies were transferred to breeding cages

Adult butterflies were kept in stock breeding cages (1 x 1.5 x 2 m) separated by species in a greenhouse with natural light. Butterflies were fed *ad libitum* using feeders containing artificial nectar made of 10% (m/v) sucrose and 1.5% (m/v) Critical Care Formula (Vetark). Breeding cages were also supplied with flowering *Lantana sp.* and *Psiguria sp.*, as a source of pollen, and with *Passiflora* plants for egg-laying.

### Morphology of plant trichomes and caterpillar prolegs

Fully unfolded leaves of *P. adenopoda* were cut off and kept fresh by placing the leaf stalks in a water-filled vial. Circular leaf discs (8 mm diameter) were cut using a custom-made hole puncher and used for either scanning electron microscopy (SEM) and electron dispersive X-ray spectroscopy (EDX) to determine ultrastructure and elemental composition of the trichomes, respectively.

The density of trichomes on *P. adenopoda* leaves was studied by placing leaf discs (8 mm diameter; n=5 from 5 different plants) under a Leica MZ16 stereomicroscope (Leica Microsystems, Wetzlar, Germany). The number of trichomes was counted on both the upper and lower leaf surfaces. The size distribution, density and orientation of the trichomes was examined using five leaves, each from different plants, under a stereomicroscope with the light source positioned perpendicular to the leaf blade. As the trichomes are translucent and difficult to see, their visibility was improved by lightly spraying some talcum powder onto the surface of the leaf (46). The leaf surfaces were viewed either at ∼ 45 degree angle (for estimating trichome density) or parallel to the microscope objective (for looking at size distribution).

For scanning electron microscopy (SEM), six leaf discs (two per species) were fixed overnight in a mixture of 4% formaldehyde and 70% ethanol, followed by dehydration in a series of increasing ethanol concentrations (70%, 80%, 90 % and absolute). The leaf discs were then critical point dried using a Quorum E3100 critical point dryer, sputter-coated with 35 nm of gold (Quorum K575X, Quorum Technologies Ltd., East Grinstead, UK) and viewed at 2 keV either with a FEI Verios 460 scanning electron microscope (FEI Company, Hillsboro, OR USA), or a Tescan MIRA3 FEG scanning electron microscope (TESCAN UK Ltd., Cambridge, UK). For EDX, two leaf discs (one disc per side) were fixed and critical point dried as for the SEM preparation. The dried leaf discs were coated with 30 nm of carbon and EDX was performed using an AMETEK window-less silicone drift detector and Genesis software. The accelerating voltage for EDX was 20 keV with a probe current of 0.4 nA and images were acquired using the concentric backscatter detector. All SEM and EDX images were analysed using Fiji (71).

To observe the morphology of the caterpillars’ prolegs, ethanol-fixed and critical point dried specimens of fourth instar caterpillars of *H. erato*, *H. melpomene*, *H. charithonia* and *D. iulia* were sputter-coated with a 35 nm layer of gold and imaged by SEM as above.

### Observation of trichome-caterpillar interaction

To analyse caterpillar-trichome interactions, experiments were performed using 3^rd^ - 4^th^ instar caterpillars of *H. melpomene, H. charithonia*, *D. iulia*, and *H. erato* on *P. adenopoda* leaves with and without trichomes (n=10 for each species and treatment). Leaves were "shaved" by using a beard trimmer (Phillips Ltd., Netherlands) to remove the trichome tips, followed by a twin-blade razor (Boots Ltd., Nottingham, UK) to remove the remaining parts of the trichome. Shaved leaves were carefully examined under a stereo-binocular microscope to check for any intact trichomes.

Caterpillar walking behaviour on *P. adenopoda* leaves was observed for 10 minutes and video recorded from above using a Nikon D750 DSLR camera at 23.98 or 59.98 frames per second. If the caterpillar left the leaf surface, it was placed back into the center of the leaf; the frequency with which this happened was recorded. The body mass of each caterpillar was measured to the nearest 0.1 mg using a laboratory balance (Sartorius Ac210s or Sartorius MC5, Sartorius AG, Göttingen, Germany).

The velocity with which caterpillars moved on leaves with or without trichomes was recorded. Tracking markers (black dots of ca. 0.1mm diameter on a small piece of white paper) were attached to the head, between the two head spikes, and to the dorsal midline of the third or fourth abdominal segment of the caterpillar, using dental wax (EliteHD+ light body, Zhermack, Badia Polesine, Italy). The caterpillar’s velocity measured using Fiji (71) from a randomly selected video section with a linear stereotyped motion of the caterpillar.

For higher-magnification video recordings, 1^st^/2^nd^ instars (10-30mg) and 3^rd^/4^th^ instar larvae (150-400mg) of the four Heliconiini species(n=5 for each instar and species) were placed individually on *P. adenopoda* leaves with trichomes and filmed using a DMK 23UP1300 camera (Imaging Source Europe GmbH, Bremen, Germany) and an Optem zoom 125 lens (Excelitas Tech. Corp., Mississauga, Canada) at 30 frames per second.

### Visualisation of cuticle injuries using methylene blue staining

Injuries on the proleg cuticle of 3^rd^/4^th^ instar caterpillars were visualized by staining with a 0.1 % solution of methylene blue in H_2_O (72, 73). Ten fourth instar larvae of each of the four Heliconiini species were stained after walking for 30 minutes on a *P. adenopoda* leaf with intact trichomes enclosed in a square Petri-dish (120 mm x 120 mm) lined with wet tissue paper. The caterpillar was then removed and placed for 1 minute in a 90 mm diameter petri-dish containing 30 ml of 0.1 % methylene blue. The amount of methylene blue was just enough to cover the prolegs, but did not harm the caterpillar. The caterpillar was then placed in a petri-dish containing water to wash away the excess stain. The caterpillar was anaesthetised by cooling at 4° C for 10 min and its ventral side was examined under a stereomicroscope and checked for blue spots indicating injuries (73). Pictures were taken using a camera (Point Grey Research Inc., British Columbia, Canada) mounted on the stereomicroscope and the total number of injuries was counted. The same experiment was repeated with 3^rd^/4^th^ instars of *H. erato*, *H. charithonia* and *D. iulia* (n=5 for each) on "shaved" *P. adenopoda* leaves. 5 caterpillars that had been subjected to the trichomes were randomly chosen and their prolegs examined under the SEM at the location of the blue dots.

### Penetrometry of caterpillar cuticle

We measured the puncture resistance of the caterpillars’ body wall and prolegs to assess the effect of hooked trichomes of *P. adenopoda* on the caterpillars’ cuticle. We used a custom-built penetrometer that has been described in a previous study (62). In brief, if consists of a fiber-optic 1D-force transducer with a 65 × 12 × 0.15 mm (free length × width × thickness) spring steel beam fixed at both ends. An insect pin mounted in the centre of the beam using the base of a shortened syringe needle as a holder. On the back side of the beam, a small piece of reflective foil served as the target for a fiber optic sensor (D12, Philtec, INC., Annapolis, USA) mounted above the beam, which measured the distance to the reflective foil (and hence the deflection of the beam) as a voltage signal. The sensor’s position could be controlled with a small 1-axis translation stage (Newport M-DS25). The force sensor was calibrated with weights (five weights, increasing from 1mg to 5 mg) to convert the voltage into a force (in mN). The fiber-optic sensor was operated in the linear range of its sensitive near field, with a standoff distance of 80 *μ*m between the sensor tip and the reflective foil target. The beam had a noise level of ca. 0.3 mN. A linear actuator (Physik Instrumente M222.20) moved the force tester with attached insect pin into and out of the sample with a constant speed of 0.61 mm s^−1^. When the pin penetrated the cuticle, the forces initially increased and then dropped rapidly when the cuticle fractured. The movement was stopped manually when the maximum had been reached; examination of the samples after the experiments showed that the cuticle samples had been punctured. We recorded the difference between the maximum deflection and the baseline to calculate the puncture force.

Penetrometry was performed on the body wall cuticle and prolegs of 3rd - 4th instar caterpillars, and on the body wall cuticle of 1st-2nd instar caterpillars of *H. erato*, *H. melpomene*, *H. charithonia* and D*. iulia*. An aluminium sample holder with a central circular hole of 0.5 mm diameter was used to clamp the cuticle. We used size 0-1 stainless steel Austerlitz insect pins (minuten pins) which are approximately conical with a rounded (hemispherical) tip. The tip radius of the pins was ca. 5 microns, approximately one order of magnitude higher than the tip radius of *P. adenopoda* trichomes. Despite this difference, the setup can quantify the variation in puncture resistance between the different species. Each pin was inspected under a light microscope and pins with irregular or blunter tips (with tip radii >5µm) were not used.

### Cuticular thickness measurements

*a) Area-specific mass.* Cuticle sections of the lateral body wall of 1^st^-2^nd^ and 4^th^ instar caterpillars and of the proleg subcorona of 4^th^ instar caterpillars were dissected out using micro-scissors in 0.1% (w/v) PBS and attached tissues (fat bodies and epidermis) were removed from the samples with forceps under a Leica MZ16 dissecting microscope. The surface area of the samples was measured from images of the sections on a 1mm x 1mm grid paper using ImageJ (74).The cuticle sections were then air-dried for 7 days and weighed using a Mettler Toledo MX5 microgram balance (Mettler-Toledo Inc., Switzerland). The area-specific mass of the sections was calculated by dividing the mass of each section by its surface area.
*b) SEM of freeze fractures.* Sections of the lateral body wall of 1^st^-2^nd^ caterpillars (∼10-15 samples per species) were fractured by quickly freezing samples in liquid nitrogen and fracturing them using a scalpel. The fractured samples were then mounted on SEM stubs using dental wax. The samples were sputter-coated with 10 nm of platinum (Quorum K575X, Quorum Technologies Ltd., East Grinstead, UK) and imaged under the Tescan MIRA3 FEG-Scanning electron microscope (TESCAN UK Ltd., Cambridge, UK), at 5eV.

### Caterpillar growth rate and survival on *P. biflora* versus *P. adenopoda*

Caterpillars of the four different Heliconiini species were reared on either *P. biflora* (subgenus *Decaloba*, no trichomes) or *P. adenopoda* (subgenus *Decaloba*, hooked trichomes) to measure their growth and survivorship. Three replicates of 30-50 fresh eggs of each species were placed on young shoot tips of *P. biflora* or *P. adenopoda* and kept in separate rearing cages. The eggs were allowed to hatch and the larvae were supplied with fresh leaves every other day to feed *ad libitum* on their food plants until they reached pupation. Pupae were collected and their weight and day of pupation recorded.

### Data analysis

Data were statistically analysed using R-4.4.1 (R Core Team, 2017).

## Acknowledgments

We would like to thank Dr Karin Müller (Cambridge Advanced Imaging Centre) and Dr Heather Greer (Department of Chemistry, University of Cambridge) for assistance with electron microscopy and Mr James Rolfe (Department of Earth Sciences, University of Cambridge) for providing access to the microgram balance. R.C. was funded by R. O. Whyte PhD Studentship and the Trinity Henry Barlow Studentship from the University of Cambridge and S.C. was supported by a Doctoral Training Partnership from the Biotechnology and Biological Sciences Research Council, University of Cambridge. E.C.P.d.C., G.J and C.D.J would like to thank the financial support of the Natural Environment Research Council grant NE/W005131/1, and W.F. would like to acknowledge the support of the Human Frontier Science Program Research Grant RGP008/2023-101.

## Author Contributions

Conceptualization: R.C., E.C.P.d.C., C.D.J., S.C. & W.F.; Methodology, Investigation and Visualization: RC; S.C. built the penetrometer; G.J. & R.C. involved in plant and insect care; Supervision: W.F., E.C.P.d.C. & C.D.J.; Writing – original draft: R.C.; Writing – review & editing: R.C., E.C.P.d.C., G.J., S.C., C.D.J. & W.F.

