## Supplementary material for "Caterpillar counter-adaptations to hooked trichomes: does the coevolutionary arms race continue?": Table 1

| Species | Survival on *P. biflora* | Survival on *P. adenopoda* |
|---|---|---|
| *H. erato* | 0.54 (n=81/151) | 0.0 (n=0/152) |
| *H. melpomene* | 0.24 (n=36/151) | 0.0 (n=0/134) |
| *H. charithonia* | 0.66 (n=82/124) | 0.19 (n=31/167) |
| *D. iulia* | 0.53 (n=80/152) | 0.0 (n=0/130) |

**Table 1.** Larval survival (number of larvae that reached pupation per number of eggs from for each plant species) on *P. adenopoda* vs *P. biflora*
