## Supplementary Materials for "Caterpillar counter-adaptations to hooked trichomes: does the coevolutionary arms race continue?"

**Supplementary Figures:**

Trichomes were measured and counted from cross-sections of five fully expanded young leaves from three different plants (one cross-section per leaf) under a light microscope. Trichome heights on both the leaf upper side and underside were measured, and most of the trichomes ranged in height from 50 to 300 μm. Trichome were larger on the upper side than on the underside (median heights: upperside=164 μm , underside=123 μm ; Wilcoxon rank sum test; n= 383; W= 7560; $p<0.001$).

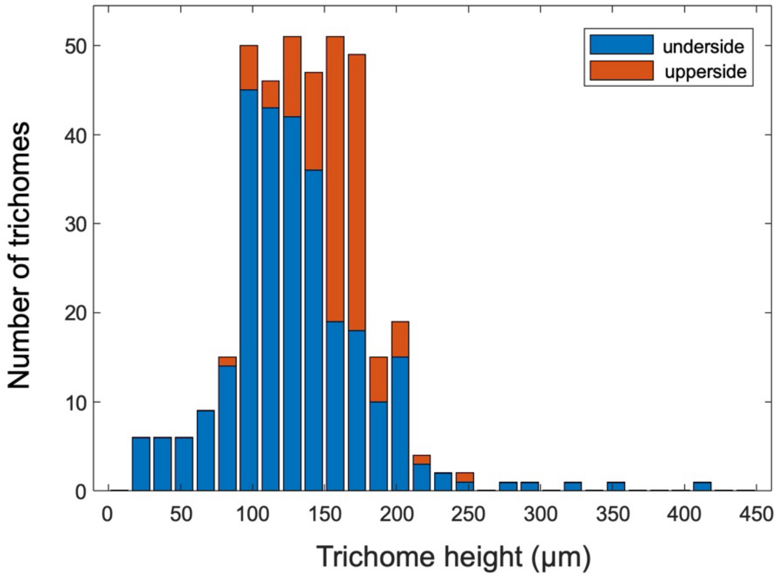


**Fig. S1.** Distribution of trichome sizes on the surface of the leaves

**Supplementary Tables:**

**Table S1.** Comparison of walking velocity of 4th instar Heliconiini caterpillars on hooked trichome-bearing (intact) leaves and "shaved" leaves. (a) Two-way ANOVA: the interaction Species x Treatment was significant, so that pairwise tests were conducted. (b) Pairwise Tukey HSD tests comparing different species for intact leaves. (c) Pairwise Tukey HSD tests comparing different species for shaved leaves. (d) Independent t-tests comparing walking velocity between intact and shaved leaves for each caterpillar species.

a) Two-way ANOVA

| | Sum of Sq. | Df | Mean Sq. | F-value | Pr(>F) |
|---|---|---|---|---|---|
| Species | 72.71 | 3 | 24.24 | 17.12 | $1.0 \times 10^{-5}$ |
| Treatment | 84.78 | 1 | 84.78 | 59.89 | $1.0 \times 10^{-5}$ |
| Interaction Species x Treatment | 24.74 | 3 | 8.25 | 5.83 | 0.0013 |
| Residuals | 94.84 | 67 | 1.42 | | |

b) Pairwise Tukey HSD tests comparing different species for intact leaves. "lwr" and "upr" indicate lower and upper bounds of confidence interval of difference.

| Species comparison | difference (mm/s) | lwr (mm/s) | upr (mm/s) | $p_{adj}$ |
|---|---|---|---|---|
| *H. charithonia - D. iulia* | -2.171 | -3.835 | -0.506 | 0.0014 |
| *H. erato - D. iulia* | -3.667 | -5.332 | -2.003 | $1.0 \times 10^{-5}$ |
| *H. melpomene - D. iulia* | -4.111 | -6.033 | -2.188 | $1.0 \times 10^{-5}$ |
| *H. erato - H. charithonia* | -1.497 | -3.161 | 0.168 | 0.109 |
| *H. melpomene - H. charithonia* | -1.940 | -3.862 | -0.018 | 0.046 |
| *H. melpomene - H. erato* | -0.443 | -2.366 | 1.479 | 0.996 |

c) Pairwise Tukey HSD tests comparing different species for shaved leaves

| Species comparison | diff (mm/s) | lwr (mm/s) | upr (mm/s) | $p_{adj}$ |
|---|---|---|---|---|
| *H. charithonia - D. iulia* | -1.221 | -2.886 | 0.443 | 0.311 |
| *H. erato - D. iulia* | -0.949 | -2.614 | 0.715 | 0.633 |

| | | | | |
|---|---|---|---|---|
| *H. melpomene* - D. iulia | -1.486 | -3.196 | 0.224 | 0.135 |
| *H. erato* - *H. charithonia* | 0.272 | -1.392 | 1.937 | 0.999 |
| *H. melpomene* - *H. charithonia* | -0.264 | -1.975 | 1.446 | 0.999 |
| *H. melpomene* - *H. erato* | -0.537 | -2.247 | 1.174 | 0.975 |

d) Independent t-tests comparing walking velocity between intact and shaved leaves within each species.

| Comparison Intact vs. Shaved | t | df | p-value |
|---|---|---|---|
| *H. charithonia* (log-transformed) | -4.73 | 21.57 | $1.1 \times 10^{-4}$ |
| *D. iulia* (log-transformed) | -1.58 | 13.89 | 0.136 |
| *H. erato* (log-transformed) | -5.83 | 15.32 | $3.0 \times 10^{-5}$ |
| *H. melpomene* (square root-transformed) | -5.06 | 8.24 | $8.9 \times 10^{-4}$ |

**Table S2.** Comparison of the number of injuries on the planta of the prolegs of 4th instar Heliconiini caterpillars after walking on *P. adenopoda* leaves. Species differed highly significantly (Kruskal-Wallis, $\chi^2 = 28.1$ df=3, p<0.001); pairwise differences were tested with Dunn's tests (with Holm correction).

| Species comparison | Z | $p_{adj}$ |
|---|---:|---:|
| *H. charithonia-D. iulia* | 1.323 | 0.371 |
| *H. erato-D. iulia* | -3.075 | 0.0084 |
| *H. melpomene-D. iulia* | -4.399 | $6.5 \times 10^{-5}$ |
| *H. erato-H. charithonia* | -2.909 | 0.0185 |
| *H. melpomene-H. charithonia* | -4.056 | $2.5 \times 10^{-4}$ |
| *H. melpomene-H. erato* | -0.246 | 0.805 |

**Table S3.** Comparison of the puncture resistance of the body wall of 1st-2nd instar Heliconiini caterpillars (penetrometry measurements). Species differed highly significantly (ANOVA, $F_{3,32}$=8.67, p<0.001); pairwise differences were tested with Tukey HSD tests.

| Species comparison | diff (mN) | lwr (mN) | upr (mN) | $p_{adj}$ |
|---|---|---|---|---|
| *H. charithonia-D. iulia* | 0.312 | -0.481 | 1.105 | 0.712 |
| *H. erato-D. iulia* | -0.812 | -1.600 | -0.019 | 0.043 |
| *H. melpomene-D. iulia* | -0.934 | -1.720 | -0.141 | 0.016 |
| *H. erato-H. charithonia* | -1.124 | -1.910 | -0.331 | 0.0029 |
| *H. melpomene-H. charithonia* | -1.246 | -2.040 | -0.453 | $9.3 \times 10^{-4}$ |
| *H. melpomene-H. erato* | -0.122 | -0.916 | 0.671 | 0.975 |

**Table S4.** Comparison of the puncture resistance of the soft planta of the proleg of 4th instar Heliconiini caterpillars (penetrometry measurements). Species differed highly significantly (ANOVA, $F_{3,27}$=10.83, p<0.001); pairwise differences were tested with Tukey HSD tests.

| Species comparison | diff (mN) | lwr (mN) | upr (mN) | $p_{adj}$ |
|---|---|---|---|---|
| *H. charithonia-D. iulia* | -0.533 | -1.622 | -0.556 | 0.546 |
| *H. erato-D. iulia* | -1.622 | -2.711 | -0.533 | 0.0019 |
| *H. melpomene-D. iulia* | -1.937 | -2.987 | -0.887 | $1.5 \times 10^{-5}$ |
| *H. erato-H. charithonia* | -1.089 | -2.244 | 0.066 | 0.069 |
| *H. melpomene-H. charithonia* | -1.404 | -2.522 | -0.285 | 0.0098 |
| *H. melpomene-H. erato* | -0.315 | -1.433 | 0.804 | 0.867 |

**Table S5.** Comparison of the puncture resistance of the body wall of 4th instar Heliconiini caterpillars (penetrometry measurements). Species differed highly significantly (ANOVA, $F_{3,36}$=9.47, p<0.001); pairwise differences were tested with Tukey HSD tests.

| Species comparison | diff (mN) | lwr (mN) | upr (mN) | $p_{adj}$ |
|---|---|---|---|---|
| *H. charithonia-D. iulia* | -1.690 | -2.743 | -0.638 | $6.3 \times 10^{-4}$ |
| *H. erato-D. iulia* | -1.889 | -2.941 | -0.837 | $1.4 \times 10^{-4}$ |
| *H. melpomene-D. iulia* | -1.321 | -2.373 | -0.269 | 0.0091 |
| *H. erato-H. charithonia* | -0.199 | -1.251 | 0.853 | 0.956 |
| *H. melpomene-H. charithonia* | 0.369 | -0.683 | 1.421 | 0.781 |
| *H. melpomene-H. erato* | 0.568 | -0.484 | 1.620 | 0.475 |

**Table S6.** Comparison of area-specific mass of the soft planta of the prolegs of 4th instar Heliconiini caterpillars. Species differed highly significantly (ANOVA, $F_{3,72}$=19.6, p<0.001); pairwise differences were tested with Tukey HSD tests.

| Species comparison | diff (µg/mm$^2$) | lwr (µg/mm$^2$) | upr (µg/mm$^2$) | $p_{adj}$ |
|---|---|---|---|---|
| *H. charithonia-D. iulia* | -0.064 | -0.469 | 0.342 | 0.976 |
| *H. erato-D. iulia* | -0.530 | -0.949 | -0.112 | 0.0072 |
| *H. melpomene-D. iulia* | -1.079 | -1.490 | -0.667 | 1.0 x $10^{-5}$ |
| *H. erato-H. charithonia* | -0.467 | -0.899 | -0.034 | 0.029 |
| *H. melpomene-H. charithonia* | -1.015 | -1.441 | -0.589 | 1.0 x $10^{-5}$ |
| *H. melpomene-H. erato* | -0.548 | -0.986 | -0.109 | 0.0082 |

**Table S7.** Comparison of area-specific mass of the body wall of 4th instar Heliconiini caterpillars. Species differed highly significantly (ANOVA, $F_{3,117}$=9.22, p<0.001); pairwise differences were tested with Tukey HSD tests.

| Species comparison | diff ($\mu g/mm^2$) | lwr ($\mu g/mm^2$) | upr ($\mu g/mm^2$) | $p_{adj}$ |
|---|---|---|---|---|
| *H. charithonia-D. iulia* | -0.559 | -0.849 | -0.269 | $1.0 \times 10^{-5}$ |
| *H. erato-D. iulia* | -0.411 | -0.704 | -0.118 | 0.0021 |
| *H. melpomene-D. iulia* | -0.369 | -0.663 | -0.077 | 0.0071 |
| *H. erato-H. charithonia* | 0.148 | -0.152 | 0.448 | 0.574 |
| *H. melpomene-H. charithonia* | 0.189 | -0.111 | 0.489 | 0.358 |
| *H. melpomene-H. erato* | 0.041 | -0.261 | 0.344 | 0.984 |

**Table S8.** Comparison of thickness of the body wall of $1^{st}$-$2^{nd}$ instar Heliconiini caterpillars (freeze-fractured samples). Species differed highly significantly (ANOVA, $F_{3,48}=23.55$, $p<0.001$); pairwise differences were tested with Tukey HSD tests.

| Species comparison | diff (µm) | lwr (µm) | upr (µm) | $p_{adj}$ |
|---|---|---|---|---|
| *H. charithonia-D. iulia* | 1.218 | -1.190 | 3.627 | 0.538 |
| *H. erato-D. iulia* | -4.292 | -6.842 | -1.742 | $2.2 \times 10^{-5}$ |
| *H. melpomene-D. iulia* | -5.645 | -8.257 | -3.034 | $1.0 \times 10^{-5}$ |
| *H. erato-H. charithonia* | -5.510 | -8.020 | -2.999 | $1.0 \times 10^{-5}$ |
| *H. melpomene-H. charithonia* | -6.864 | -9.437 | -4.291 | $1.0 \times 10^{-5}$ |
| *H. melpomene-H. erato* | -1.354 | -4.059 | 1.352 | 0.547 |

**Movie S1 (separate file).** 2nd instar *Heliconius erato* caterpillar being injured by *Passiflora adenopoda* trichomes

**Movie S2 (separate file).** Soft planta of the prolegs of 4th instar *H. melpomene* caterpillar being injured by *P. adenopoda* trichomes

**Movie S3 (separate file).** 4th instar *H. charithonia* caterpillar walking over hooked trichomes on *P. adenopoda* without being injured

**Movie S4 (separate file).** 4th instar *Dryas iulia* caterpillar walking over hooked trichomes on *P. adenopoda* without being injured
